# Regulation of hair cell and stomatal size by a hair-cell specific peroxidase in the grass *Brachypodium distachyon*

**DOI:** 10.1101/2022.07.03.498611

**Authors:** Tiago D. G. Nunes, Lea S. Berg, Magdalena W. Slawinska, Dan Zhang, Leonie Redt, Richard Sibout, John P. Vogel, Debbie Laudencia-Chingcuanco, Barbara Jesenofsky, Heike Lindner, Michael T. Raissig

## Abstract

The leaf epidermis is the outermost cell layer forming the interface between plants and the atmosphere that must both provide a robust barrier against (a)biotic stressors and facilitate carbon dioxide uptake and leaf transpiration ^1^. To achieve these opposing requirements, the plant epidermis developed a wide range of specialized cell types such as stomata and hair cells. While factors forming these individual cell types are known ^2–5^, it is poorly understood how their number and size is coordinated. Here, we identified a role for *BdPRX76*/*BdPOX*, a class III peroxidase, in regulating hair cell and stomatal size in the model grass *Brachypodium distachyon*. In *bdpox* mutants prickle hair cells were smaller and stomata were longer. Because stomatal density remained unchanged, the negative correlation between stomatal size and density was disrupted in *bdpox* and resulted in higher stomatal conductance and lower intrinsic water-use efficiency. *BdPOX* was exclusively expressed in hair cells suggesting that *BdPOX* cell-autonomously promotes hair cell size and indirectly restricts stomatal length. Cell wall autofluorescence and lignin stainings indicated a role for BdPOX in lignification or crosslinking of related phenolic compounds at the hair cell base. Ectopic expression of *BdPOX* in the stomatal lineage increased phenolic autofluorescence in guard cell walls and restricted stomatal elongation in *bdpox*. Together, we highlight a developmental interplay between hair cells and stomata that optimizes epidermal functionality. We propose that cell-type-specific changes disrupt this interplay and lead to compensatory developmental defects in other epidermal cell types.

## RESULTS AND DISCUSSION

### *bdpox* mutants show increased stomatal conductance and cell size defects in stomata and prickle hair cells

In the model grass *B. distachyon*, the leaf blade epidermis is dominated by rectangular pavement cells, stomatal complexes, trichomes consisting of prickle hair cells (PHCs) and few interspersed macrohairs, and occasional silica cells (Fig. 1A). Most specialized amongst epidermal cells are stomatal complexes, which are cellular pores on the epidermis that drive plant gas exchange. Fast stomatal movements and adjustments in stomatal anatomy (stomatal density and size) are crucial for water-use efficient gas exchange thereby contributing to abiotic stress resilience ^6,7^. Yet, since stomatal density and size are negatively correlated, it remains elusive how modifying just a single of these anatomical traits affects stomatal function.

**Figure 1.**
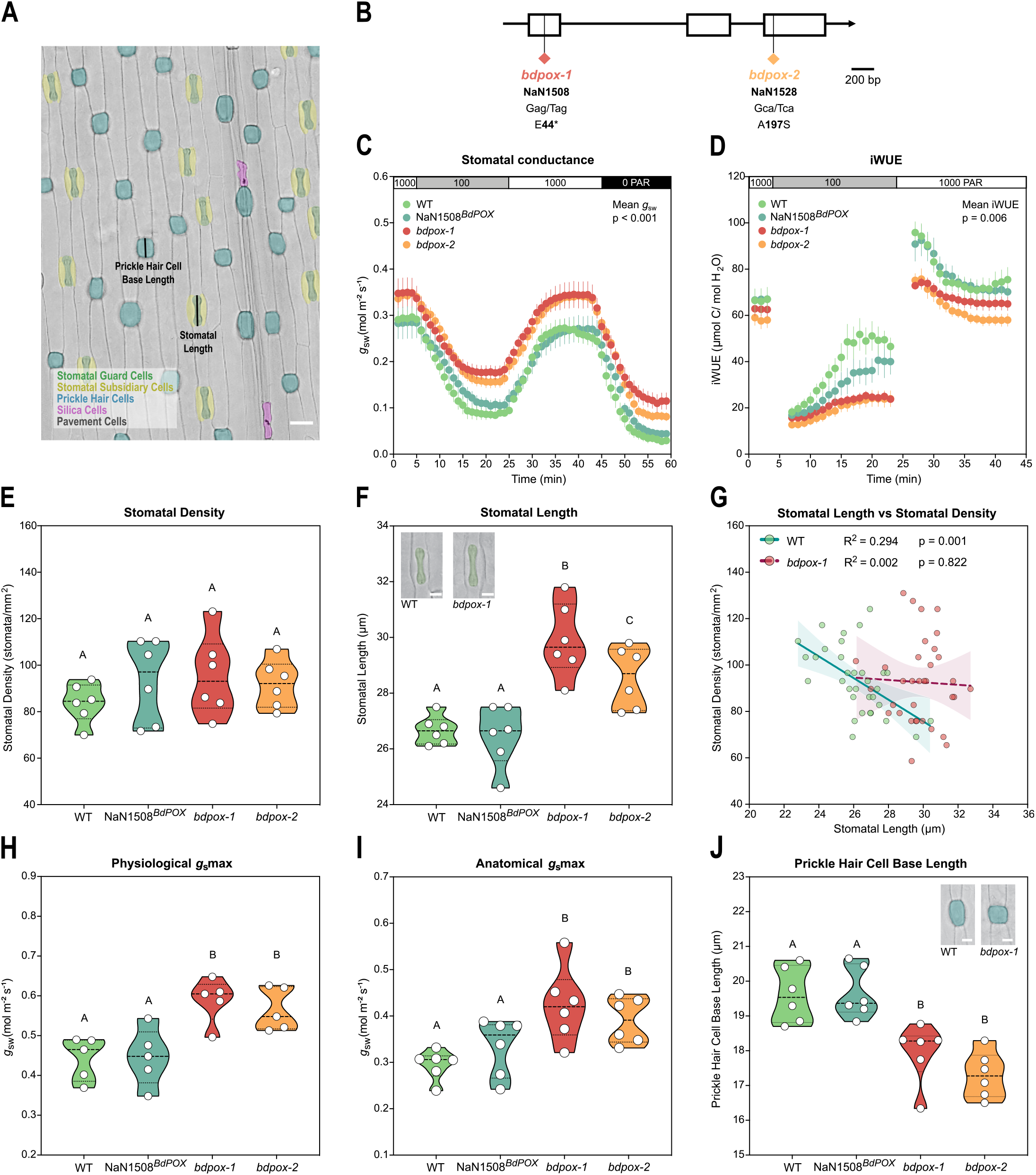
*bdpox* mutants display altered gas exchange and anatomical defects in the leaf epidermis. **(A)** Leaf epidermis of *B. distachyon* (Bd21-3); stomatal complexes (GCs in green and SCs in yellow), prickle hair cells (blue), pavement cells (grey) and silica cells (magenta). Measuring axis of stomatal length (SL) and PHC base length (PHC BL) are indicated. Scale bar, 20 μm. **(B)** *BdPRX76/BdPOX* (BdiBd21-3.2G0467800) gene model indicating the location and nature of the mutations in *bdpox-1* (NaN1508) and *bdpox-2* (NaN1528). **(C)** Stomatal conductance (*g*_sw_) in response to changing light (1000-100-1000-0 PAR) in WT, NaN1508^*BdPOX*^, *bdpox-1* and *bdpox-2* (n=6 individuals per genotype). Dots represent the mean and error bars represent SEM. p value obtained from one-way ANOVA comparing differences on mean *g*_sw_ among different groups indicated in the graph. Full statistical analysis demonstrating significant differences between wild-type and *bdpox* mutants in Supplementary dataset. **(D)** Intrinsic water-use efficiency (iWUE) in response to changing light (1000-100-1000 PAR) in WT, NaN1508^*BdPOX*^, *bdpox-1* and *bdpox-2* (n=6 individuals per genotype). Dots represent the mean and error bars represent SEM; p-value obtained from one-way ANOVA comparing differences on mean *g*_sw_ among different groups is indicated in the graph. Full statistical analysis demonstrating significant differences between wild-type and *bdpox* mutants can be found in Supplementary dataset. **(E)** Stomatal density (SD) in WT, NaN1508^*Bdpox*^, *bdpox-1* and *bdpox-2* (n=6 individuals per genotype, 548-619 stomata counted per genotype). **(F)** Stomatal length (SL) in WT, NaN1508^*Bdpox*^, *bdpox-1* and *bdpox-2* (n=6 individuals per genotype; 193-287 stomata per genotype). Inset shows a DIC image of WT and *bdpox-1* stomata; GCs are false-colored in green; scale bar, 10 μm. **(G)** Correlation between average stomatal length (SL) and average stomatal density (SD) in WT and *bdpox-1* (n=33-34 individuals per genotype grown between Oct/2019 and May/2021). Linear regressions individually performed for WT (green) and *bdpox-1* (red). 95 % confidence bands are shown for WT (green) and *bdpox-1* (red). R^2^ and p-values are indicated, dashed line represents non-significant correlation (slope not significantly different than zero). **(H)** Physiological *g*_s_max measurements in WT, NaN1508^*Bdpox*^, *bdpox-1* and *bdpox-2* (n=5 individuals per genotype). **(I)** Anatomical *g*_s_max calculations in WT, NaN1508^*Bdpox*^, *bdpox-1* and *bdpox-2* (n=6 individuals per genotype; same individuals as in C-F). **(J)** PHC base length in WT, NaN1508^*BdPOX*^, *bdpox-1* and *bdpox-2* (n=6 individuals; 656-691 PHCs per genotype). Inset shows DIC image of WT and *bdpox-1* PHCs; PHCs false-colored in blue; scale bar, 10 μm. Different letters represent significant differences (p<0.05) obtained from one-way ANOVA followed by Tukey’s multiple comparisons. Data results from two independent experiments. See also Figure S1 and Figure S2.

To identify novel factors associated with graminoid stomatal morphology and functionality, we performed RNA-sequencing of mature zones of 7 day old leaves in *B. distachyon* Bd21-3 (WT) and *bdmute* plants (Fig. S1A). The *bdmute* leaf epidermis features abnormal stomata that lack SCs, which strongly affects stomatal responsiveness and gas exchange ^3^. 179 genes were downregulated in *bdmute* (Supplementary Dataset) and we selected ~50 candidate genes for a reverse genetic screening. Candidates were chosen according to their annotated gene function, their expression being lower in the developmental zone ^8^ and the availability of mutants from a collection of sodium azide (NaN_3_) mutagenized and fully resequenced lines ^9^.

In the initial screen, we found that two mutants of the class III peroxidase *BdPRX76/BdPOX* (BdiBd21-3.2G0467800; Fig. 1B; S1A, B) showed lower intrinsic water-use efficiency (iWUE; Fig. S1C) and higher ambient-adapted stomatal conductance (*g*_sw_; Fig. S1D). The two NaN mutants disrupting *BdPRX76/BdPOX* were NaN1508, which contained a heterozygous, early STOP codon (E44*; *bdpox-1*) and NaN1528, which contained a homozygous missense mutation in the BdPOX active/heme-binding site (A197S; *bdpox-2;* Fig. 1B, S1B). From the segregating NaN1508 population, homozygous mutant individuals (*bdpox-1*, NaN1508^*bdpox-1*^) and wildtype segregants (NaN1508^*BdPOX*^) were selected by genotyping. Because NaN1508^*BdPOX*^ contained the same background mutations as *bdpox-1*, we included it as an additional wild-type control line.

To confirm the gas exchange defects in *bdpox* mutants, we measured stomatal conductance (*g*_sw_) in response to changing light intensity (1000-100-1000-0 PAR) ^7^. In *bdpox* plants, we observed higher *g*_sw_ in all light steps (Fig. 1C), but no significant impact on stomatal opening and closure speed (Fig. S2A-C). Since no significant variation in carbon assimilation (*A*) was observed (Fig. S2D), *bdpox* mutants suffered a decrease in intrinsic water-use efficiency (iWUE, *A/g_sw_*), particularly in the light-limited step (100 PAR; Fig. 1D).

To test if the increased *g_sw_* was caused by changes in stomatal density we performed microscopic analysis of the leaf epidermis from the leaves assessed for gas exchange. No differences were found regarding stomatal density (SD; Fig. 1E), yet we observed significantly longer stomata in *bdpox* mutants (Fig. 1F). This suggested that the well-established negative correlation between stomatal size and density, observed both interspecifically ^10–15^ and intraspecifically ^7,16,17^, was disrupted in *bdpox* (Fig. 1G). Detailed morphometric confocal analysis of fusicoccin-treated leaves to induce full stomatal opening revealed that stomatal pores are indeed longer and larger in *bdpox* (Fig. S2F-I).

To test whether the higher *g_sw_* levels could be explained by the disrupted stomatal anatomy in *bdpox*, we compared physiological *g*_s_max measurements with anatomical *g*_s_max calculations (theoretical *g*_s_max based on gas diffusion physical constants and stomatal anatomical traits) using an established equation recently optimized for grass stomata ^7^. Physiological *g*_s_max measurements confirmed the increased *g_sw_* capacity in *bdpox* mutants (Fig. 1H), and anatomical *g*_s_max calculations revealed the same relative variation between *bdpox* mutants and WT (Fig. 1I). Together, this strongly suggested a causal relationship between longer stomata and higher gas exchange in *bdpox*.

To verify if the cell size defect was specific to stomata, we also measured the length of pavement cells (PCs) and of the base of prickle hair cells (PHCs). While no differences were found in PC size among genotypes (Fig. S2J), we observed an unexpected decrease in the base length of PHCs in *bdpox* mutants (Fig. 1J).

In summary, *BdPOX* seemed to negatively affect stomatal size but positively regulate PHC size.

### *BdPOX* is expressed in hair cells and mutant complementation rescued stomatal and prickle hair cell phenotypes

To determine where *BdPOX* was expressed, we generated transcriptional (*BdPOXp:3xNLS-eGFP*) and translational (*BdPOXp:BdPOX-mCitrine*) reporter lines. To our surprise, both *BdPOX* reporter genes were exclusively expressed in PHCs (Fig. 2A, B; S3B, Movie S1) and macrohairs (Fig. S3A). Because grass leaf development and, consequently, epidermal development follows a strict base-to-tip developmental gradient with well-established stomatal stages, we used the stomatal stages as landmarks to track PHC development ^18^. We observed that both transcriptional (*BdPOXp:3xNLS-eGFP*) and translational (*BdPOXp:BdPOX-mCitrine*) reporter expression in PHCs started during stages 5-6i of stomatal development and, therefore, before significant stomatal elongation (Fig. 2A, B).

**Figure 2.**
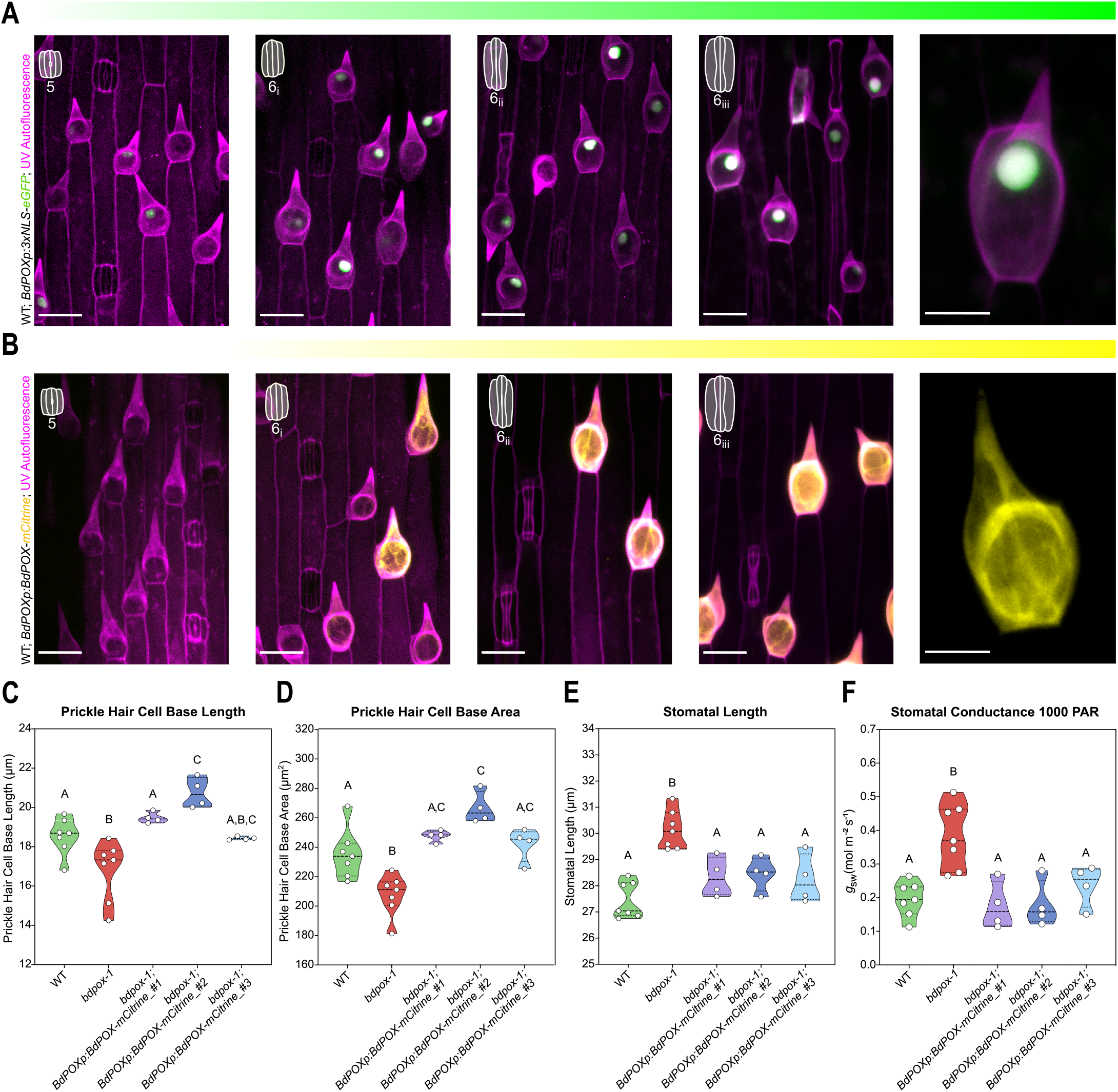
*BdPOX* reporter lines are specifically expressed in hair cells and complement the anatomical and physiological defects in *bdpox-1*. **(A)** Transcriptional reporter *BdPOXp:3xNLS-eGFP* expression stages in the developing epidermis staged according to stomatal developmental stages (upper left insets); cell wall UV-induced autofluorescence indicates cell outlines (magenta). Right-most panel shows a mature PHC. **(B)** Translational reporter *BdPOXp:BdPOX-mCitrine* expression stages in the developing epidermis staged according to stomatal developmental stages (upper left insets); cell wall UV-induced autofluorescence indicates cell outlines (magenta). Right-most panel shows a mature PHC (mCitrine channel only). **(C)** PHC base length in WT, *bdpox-1* and in three independent complementation lines (*bdpox-1; BdPOXp:BdPOX-mCitrine #1, #2, #3);* n=4-7 individuals per genotype, 160-340 PHCs per genotype/line. **(D)** PHC base area in WT, *bdpox-1* and in three independent complementation lines (*bdpox-1; BdPOXp:BdPOX-mCitrine #1, #2, #3);* n=4-7 individuals per genotype, 128-358 PHCs per genotype/line. **(E)** Stomatal length in WT, *bdpox-1* and in three independent complementation lines (*bdpox-1; BdPOXp:BdPOX-mCitrine #1, #2, #3);* n=4-7 individuals per genotype, n=156-355 stomata per genotype/line. **(F)** Steady-state stomatal conductance at 1000 PAR in WT, *bdpox-1* and in three independent complementation lines (*bdpox-1; BdPOXp:BdPOX-mCitrine #1, #2, #3);* n=4-7 individuals per genotype/line. Scale bars, 20 μm (except in right-most, mature PHC panels, where scale bar is 10 μm). Each dot represents the average of one individual. Different letters represent significant differences (p<0.05) obtained from one-way ANOVA followed by Tukey’s multiple comparisons. Data results from two independent experiments. See also Figure S3 and Movie S1.

Expression of *BdPOXp:BdPOX-mCitrine* in *bdpox-1* fully complemented both the PHC and stomatal size phenotypes (Fig. 2C-E). Detailed PHC phenotyping revealed that PHC base length, area, and outgrowth which were decreased in *bdpox-1*, were rescued in three independent *bdpox-l;BdPOX-p:BdPOX-mCitrine* complementation lines (Fig. 2C, D, S3C). Importantly, PHC base area was positively correlated with PHC outgrowth, indicating that PHC base measurements are a good proxy for PHC size (Fig. S3D). Stomatal length (SL) was rescued to WT levels in all three independent complementation lines (Fig. 2E) and stomatal density (SD) remained unaltered (Fig. S3E). Consequently, stomatal conductance (*g*_sw_) was restored to wild-type levels in the complementation lines (Fig. 2F).

Ubiquitous expression of *BdPOX* in *bdpox1*, however, had no effect on cell size nor on gas exchange suggesting that a cell type-specific expression is necessary for its complementation (Fig. S3F-I). *ZmUbip-driven* BdPOX was also expressed at significantly lower levels in PHCs (0.5 fold decrease) compared to BdPOX driven by its endogenous promoter in the *BdPOXp:BdPOX-mCitrine* line (Fig. S3F, J, K). Therefore, not only correct spatiotemporal expression but also correct dosage might be required to complement the mutant phenotype.

Together, our results suggested that *BdPOX* played a cell-autonomous role in promoting PHC size and, as a consequence, indirectly restricted stomatal elongation.

### *BdPOX* might be involved in lignification/hydroxycinnamates cross-linking at the base of prickle hair cells to positively regulate cell outgrowth

To mechanistically link how a hair-cell localized class III peroxidase could affect stomatal anatomy, we first investigated the cell-autonomous function of BdPOX in PHCs. *BdPRX76/BdPOX* seems to be in a monocot-specific clade ^19^, yet its closest *Arabidopsis* homolog, *AtPRX66*, is associated with phenolic modifications in the cell wall, namely lignification of tracheary elements ^20^. Class III peroxidases can function through a hydroxylic cycle consuming H_2_O_2_ or through a peroxidative cycle, producing H_2_O_2_ and participating in the polymerization and crosslinking of phenolic compounds (including monolignols into lignins) ^21–24^. A unique feature of grass cell walls is the significant yet cell-type-specific amount of hydroxycinnamates (ferulic acid and ρ-coumaric acids) that are bound to arabinoxylans and/or to lignins ^25–28^. Thus, we hypothesized that BdPOX modulates PHC size by altering phenolic compounds in the cell walls.

To test this, we assessed UV-induced autofluorescence of phenolic compounds in the cell wall of PHCs ^29–31^. PHC autofluorescence plot profiles and corrected total cell fluorescence revealed lower autofluorescence in *bdpox-1* compared to WT and complemented *bdpox-1*, specifically at the base of the PHCs (approximately the initial 12 μm; Fig. 3A-C, S4A).

**Figure 3.**
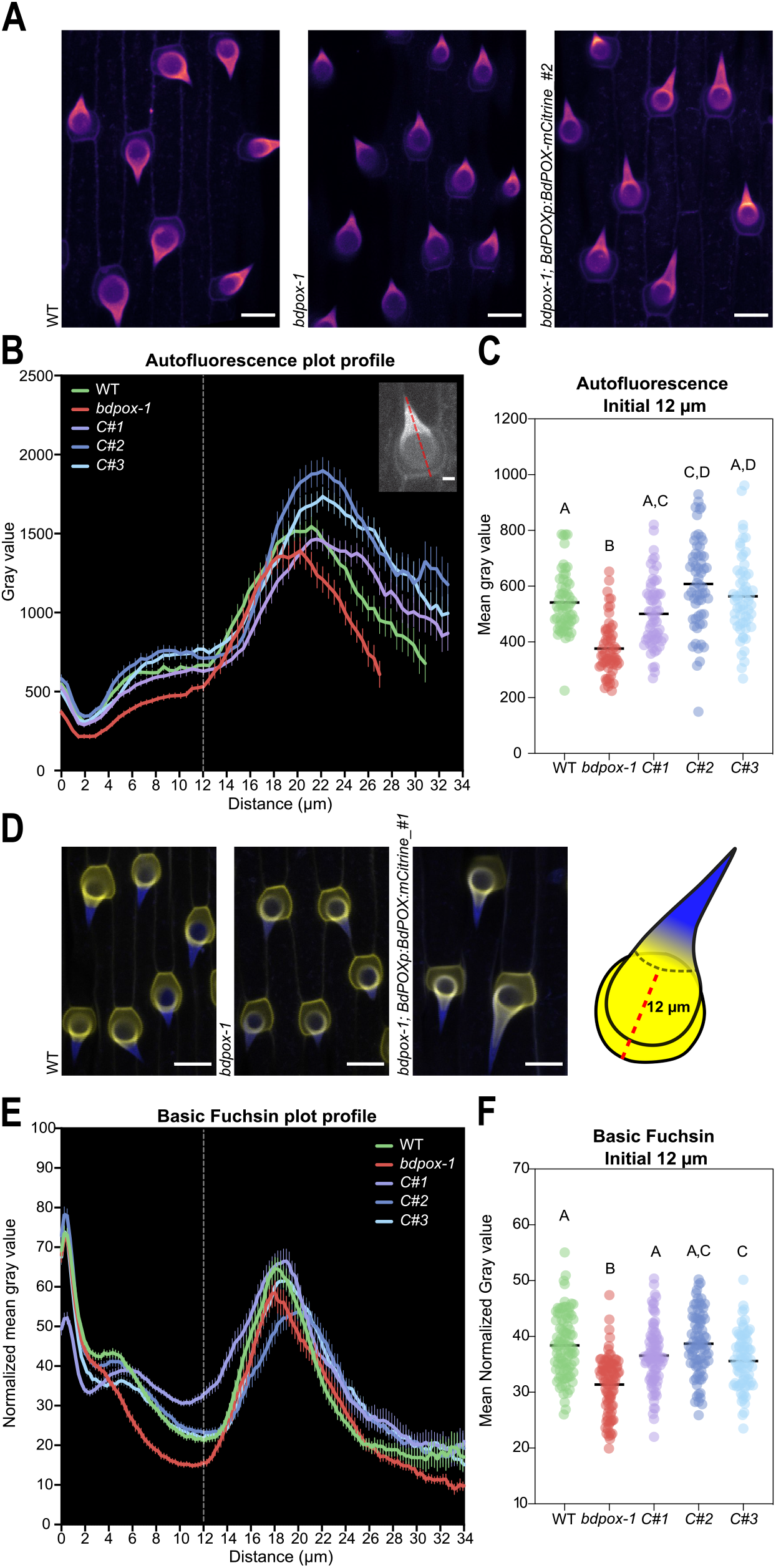
*BdPOX* affects cell wall phenolic content through lignification and/or hydroxycinnamates crosslinking. **(A)** Cell wall autofluorescence of the mature epidermis in WT, *bdpox-1* and in *bdpox-1; BdPOXp:BdPOX-mCitrine #2*. Scale bar, 20 μm. **(B)** Average plot profiles of cell wall autofluorescence in PHCs in WT, *bdpox-1* and three complementation lines (*bdpox-1; BdPOXp:BdPOX-mCitrine #1, #2, #3);* n=60 PHCs per genotype. Error bars represent SEM. Scale bar, 5 μm. **(C)** Average cell wall autofluorescence of the initial 12 micrometers of the PHCs in WT, *bdpox-1* and three complementation lines (*bdpox-1; BdPOXp:BdPOX-mCitrine #1, #2, #3);* n=60 PHCs per genotype. **(D)** Simultaneous imaging of phenolics autofluorescence (blue) and fuchsin-stained lignin (yellow) in WT, *bdpox-1* and *bdpox-1; BdPOXp:BdPOX-mCitrine #1*. Scale bars, 20 μm. **(E)** Average plot profiles of fuchsin-stained lignin in PHCs of WT, *bdpox-1* and three complementation lines (*bdpox-1; BdPOXp:BdPOX-mCitrine #1, #2, #3);* n=75-90 PHCs per genotype. Error bars represent SEM. **(F)** Basic fuchsin stain fluorescence in the initial 12 μm of the PHCs in WT, *bdpox-1* and three complementation lines (*bdpox-1; BdPOXp:BdPOX-mCitrine #1, #2, #3);* n=75-90 PHCs per genotype. Different letters represent significant differences (p<0.05) from one-way ANOVA followed by Tukey’s multiple comparisons. See also Figure S4, Movie S2 and Movie S3.

To test if the lower cell wall autofluorescence originated from decreased lignin content, we used different histochemical stainings. Basic fuchsin is a standard lignin stain ^32–34^ that also has a high affinity for hydroxycinnamates in the *B. distachyon* cell wall ^35^. Simultaneous imaging of cell wall autofluorescence and fuchsin-stained lignin showed that fuchsin preferentially stained the lower section of PHCs while total phenolics autofluorescence was observed until the tip (Fig. 3D-F; Movie S2, Movie S3). Indeed, *bdpox-1* mutants showed lower fuchsin fluorescence intensity at the basal section of PHCs (first 12 μm from the basal outline) compared to WT and complemented *bdpox-1* lines (Fig. 3E, F) suggesting reduced lignin/hydroxycinnamates content in the mutant. Very similar results were observed using safranin-O lignin staining (Fig. S4B, C), in which increased red fluorescence is observed in lignified cells, whereas non-lignified cell walls preferentially fluoresce in green ^36^. Therefore, a red/green ratiometric analysis allows for a semi-quantitative evaluation of cell wall lignification ^37^. *bdpox-1* mutants displayed a lower safranin-O ratio at the base section of PHCs (first 12 μm) compared to WT and complemented *bdpox-1*, again indicating a decrease in lignin content in the mutant (Fig. S4B, C).

Regarding stomata, no significant differences were observed in autofluorescence nor in fuchsin-stained lignin in mature GCs between WT and *bdpox-1* (Fig. S4D, E). When looking at the developing stomata during stomatal elongation/maturation, we observed that cell wall autofluorescence increased in wild-type GCs (Fig. S4F). This increase appeared to start earlier in *bdpox-1* but stalled sooner, too, which may be linked to the stomatal elongation defects in the mutants (Fig. S4F).

Overall, our data suggests that *BdPOX* participates in lignification of the basal cell wall of PHCs, which seems to be required for proper PHC growth, and indirectly impacts stomatal elongation.

### Specific expression of *BdPOX* in the stomatal lineage arrests stomatal elongation

*BdPOX* appeared to be involved in cell wall phenolic modifications (lignification/crosslinking) at the base of PHCs to promote cell outgrowth. How this PHC-specific process, however, affected stomatal elongation remained elusive. When following the basipetal developmental gradient of the *B. distachyon* epidermis, we found that PHCs grow and mature significantly before the stomatal complexes start to elongate (Fig. 4A). PHC outgrowth started when stomata were in early stages of development (i.e. stage 3-4 during SC recruitment) and was completed before GCs elongated and acquired the mature dumbbell morphology (Fig. 4A). Therefore, the growth restriction of PHCs in *bdpox* could secondarily influence stomatal anatomy, but not *vice versa*. Accordingly, the ectopic expression of *BdPOX* in the developing GC lineage using a stomatal-lineage specific promoter (*BdMUTEp:Bd-POX(CDS)-mCitrine;* Fig. 4B) significantly restricted stomatal elongation in *bdpox-1*, which correlated with increased phenolics autofluorescence in GCs (Fig. 4C, D) but not in fuchsin staining as observed in PHCs (Fig. S4G). Therefore, the ectopic expression of BdPOX in GCs seems to affect different polyphenolic compounds than lignin/hydroxycinnamates, which restricts excessive GC elongation in *bdpox* mutants (Fig. 4D). PHC size, however, remained unaffected (Fig. 4E) as PHCs mature before stomata elongate.

**Figure 4.**
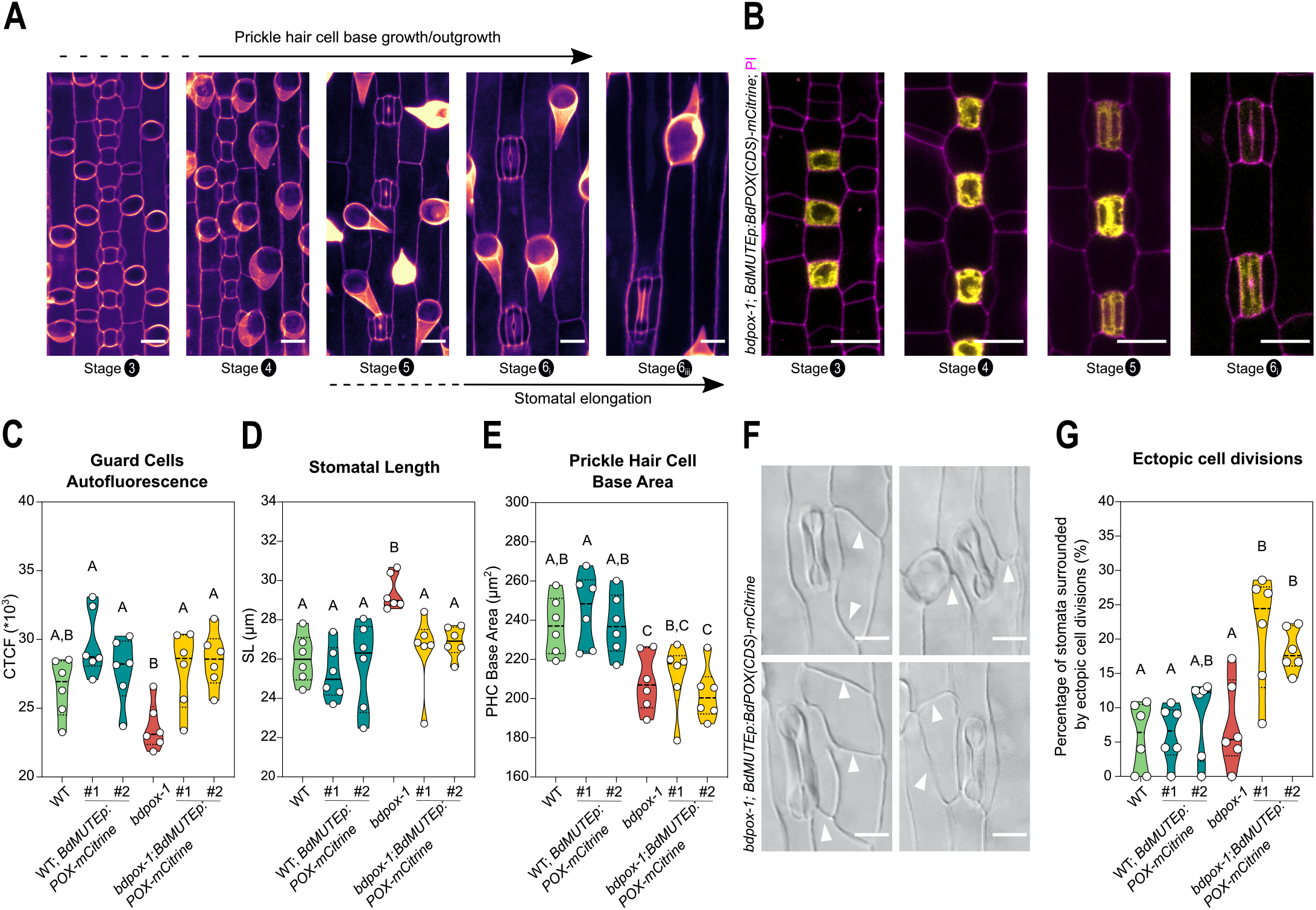
Guard cell-specific expression of *BdPOX* inhibits stomatal elongation and indicates a tissue-wide, compensatory cell elongation mechanism. **(A)** Stages of prickle hair cell development and respective stomatal development stages (stage 3 - 6). PHC differentiation, outgrowth and morphogenesis starts when the stomatal lineages are still dividing and concludes before stomatal complexes are fully elongated. Scale bars, 10 μm. **(B)** *BdMUTEp:BdPOX(CDS)-mCitrine* expression in the developing GC lineage. *BdPOX(CDS)-mCitrine* expression in GMCs (stage 3), GMCs after SC recruitment (stage 4), early developing GCs (stage 5) and early elongating GCs (stage 6i; from left to right). Scale bar, 10 μm. **(C)** Quantification of cell wall autofluorescence (CTCF, corrected total cell fluorescence) of GCs (n=6 individuals per genotype/line, 95-100 stomata per genotype,) in 2^nd^ leaves from WT, *bdpox-1* and *BdMUTEp:BdPOX(CDS)-mCitrine* lines expressed in WT and *bdpox-1*. **(D)** Quantification of stomatal length (n=6 individuals per genotype/line, 124-145 stomata per genotype/line) in mature leaves from WT, *bdpox-1* and *BdMUTEp:BdPOX(CDS)-mCitrine* lines expressed in WT and *bdpox-1*. **(E)** Quantification of PHC base area in mature leaves from WT, *bdpox-1* and *BdMUTEp:BdPOX(CDS)-mCitrine* lines expressed in WT and *bdpox-1* (n=6 individuals per genotype/line, 135-147 PHCs per genotype/line). **(F)** Stomata associated with ectopic cell divisions in the neighboring pavement cells when *BdMUTEp:BdPOX(CDS)-mCitrine* is expressed in *bdpox-1*. Scale bars, 10 μm. **(G)** Quantification of the percentage of stomata showing ectopic cell divisions in their neighboring pavement cells in mature leaves of WT, *bdpox-1* and *BdMUTEp:BdPOX(CDS)-mCitrine* expressed in WT and *bdpox-1* (n=6 individuals per genotype/line; 159-181 stomata per genotype/line). Different letters represent significant differences (p<0.05) obtained from one-way ANOVAs followed by Tukey’s multiple comparisons. See also Figure S4.

Intriguingly, we observed aberrant cell divisions in the pavement cells surrounding stomata when expressing *Bd-MUTEp:BdPOX(CDS)-mCitrine* in *bdpox-1* (Fig. 4F, G). This suggested that elongating stomata might compensate for tissue-wide mechanical imbalances caused by too small PHCs in the *bdpox-1* epidermis. Thus, when ectopic, stomatal lineage-specific expression of *BdPOX* inhibited the compensatory stomatal elongation in *bdpox-1*, additional divisions in pavement cells around stomata might be triggered to compensate for these tissue-wide mechanical tensions instead.

In conclusion, ectopically expressed *BdPOX* in GCs arrested the excessive stomatal elongation in *bdpox-1*, potentially by modifying phenolic cell wall components distinct from lignin/hydroxycinnamates. Furthermore, the observation of ectopic pavement cell divisions around length-restricted stomata in *bdpox* suggests that a growth disruption in one epidermal cell-type leads to compensatory developmental defects in other epidermal cells.

## Conclusions

The leaf epidermis is the barrier between the inner photosynthetically active tissues and the environment. Highly specialized epidermal cell types like stomata and hair cells facilitate the contrasting functional requirements of this outermost barrier. Several studies suggest that hair cell patterning intersects with the core stomatal developmental programs in *Arabidopsis ^38–41^*. Also in grasses, failing to specify stomatal identity results in hair cells being formed in their place in *B. distachyon* ^2^ and failing to specify hair cell identity leads to ectopic stomata in maize ^5^. This suggests that these cell types are ontogenetically closely related in grasses and that their development is thus likely coordinated. Furthermore, stomatal and trichome densities were shown to be negatively correlated in *Solanum lycopersicum*. The stomata to trichome ratio determined water-use efficiency ^42^ suggesting a physiological relevance for the coordination between the two cell types. Yet, the mechanisms that coordinate the formation and growth of stomata and trichomes remain highly unexplored.

Many core players guiding grass stomatal development were characterized ^2,3,18,43–50^, whereas grass hair cell formation remains poorly explored ^41^. Few grass trichome initiation factors such as the transcription factors HAIRY LEAF 6 (HL6) ^51–53^ and SQUAMOSA PROMOTER BINDING PROTEIN-LIKE 10/14/26 (SPL10/14/26) ^5,54,55^ were identified in rice and maize, but factors that affect morphogenesis and size remain mostly unknown. Here, we identified *BdPOX* and revealed its role in regulating PHC size, which indirectly affects stomatal size and optimal water-use efficiency in the model grass *B. distachyon. BdPOX* was exclusively expressed in the hair cells and seemed to participate in the lignification or/ and cross-linking of cell wall phenolics (such as hydroxycinnamates) specifically at the base of PHCs. Since lignin is a cell wall polymer that provides mechanical support ^56–58^ we speculate that such cell wall modifications at the base of PHCs are required to increase tensile strength and provide physical support for cell outgrowth. Similar processes are required in tip-growing root hairs and pollen tubes. In both cases, mechanical anisotropy along the main growth axis, which is mediated by modifying cell wall properties, seems to support cell outgrowth ^59–61^. Sustained perpendicular PHC outgrow might then feedback on PHC base expansion to maintain optimal geometrical proportions of these cells.

When BdPOX was misexpressed in *bdpox* GCs, cellular growth was restricted rather than promoted. This differential effect on cell elongation likely has several reasons; first, different polyphenolic compounds were affected in the GC context compared to the PHC context. This might have distinct effects on cell wall properties and cellular mechanics. Second and unlike in PHCs, ectopic *BdPOX* expression in the GC context occurs much before cell elongation commences potentially leading to premature cell wall stiffening and, thus, growth restriction. Third, PHCs grow perpendicular rather than parallel to the principal direction of leaf growth like GCs. Consequently, (localized) modification of cell wall properties might affect growth differently in tip-growing versus non-tip-growing cells.

The reduction in PHC size in *bdpox* indirectly altered stomatal size, but did not translate to changes in stomatal density. The resulting disruption of the negative correlation between stomatal size and density likely has two reasons; first, changes to stomatal size occur much later in development than the determination of stomatal density. Thus, changes in stomatal density can affect stomatal size *a posteriori*, where an increase in stomatal numbers can induce a downstream effect on the cell-wall machinery controlling stomatal elongation. This process is very unlikely to happen in the other direction particularly in grasses, where early stages are not only temporally but also spatially separated from late stages. Second, *bdpox* primarily impacts the PHC lineage and only affects stomatal development as a secondary consequence. Without a disruption of the stomatal genetic toolbox itself a compensatory mechanism might not be induced in a timely manner.

However, the exact mechanism of how restricting PHC growth induces stomatal elongation remains vague. We speculate that decreased PHC size may lead to changes in mechanical and/or geometrical constraints in the epidermal tissue, which would allow for increased stomatal elongation as a compensatory mechanism to reconstitute the tensile balance in the epidermis. The increase in stomatal length observed in *bdpox* mutants (~9 %) was quantitatively equivalent to the decrease in PHC base length (~10 %). In addition, expressing *BdPOX* in the GC lineage of *bdpox-1* resulted in an epidermis containing both shorter stomata and shorter PHCs and induced aberrant cell divisions surrounding the stomatal complexes. We speculate that the combination of restricted PHC growth due to *bdpox-1* and a prevention of stomatal elongation due to GC-expressed *BdPOX* may have caused a mechanical imbalance in the elongating epidermis resulting in cell divisions to compensate cellular tensions particularly around stomatal complexes. Alternatively, changes in hydrogen peroxide levels in the PHC apoplast due to loss of *BdPOX* might affect the reactive oxygen species signaling landscape, which could influence stomatal length non-cell-autonomously.

Regardless, the unique disruption in the negative correlation between stomatal size and density allowed us to investigate how modifying a single stomatal anatomical trait (i.e. stomatal size) would affect gas exchange. While an increase in stomatal size enhanced stomatal conductance, it did not significantly affect stomatal opening and closing speed, corroborating our previous observation that stomatal speed was correlated with stomatal density but not with stomatal size in *B. distachyon* ^7^.

Overall, we identified a hair cell-specific factor and demonstrated how a cell-type-specific disruption of PHC growth indirectly affected stomatal development. Strikingly, this indirect effect allowed for the specific manipulation of stomatal size without affecting stomatal density. This enabled us to test how the manipulation of a single stomatal anatomy trait (i.e. stomatal size) affected stomatal gas exchange physiology and water-use efficiency in a grass model for the first time. Consequently, manipulating PHC size might present an indirect route to potentially alter stomatal size in grasses without affecting stomatal density.

## ACKNOWLEDGMENTS

The authors thank Prof. Dr. Annika Guse and Prof. Dr. Jochen Wittbrodt for access to microscope facilities, Prof. Dr. Karin Schumacher, Prof. Dr. Rüdiger Hell, Dr. Graham Dow, Dr. Upendo Lupanda, and Dr. Paula Ragel for scientific and technical insights. We thank Michael Schilbach for greenhouse facility managment and gardening support. We also acknowledge Prof. Dr. Dominique Bergmann and the Howard Hughes Medical Institute for supporting the RNA-sequencing experiment and Laura R. Lee for help with RNA-seq data analysis. The work (proposal: 10.46936/10.25585/60001041) conducted by the U.S. Department of Energy Joint Genome Institute (https://ror.org/04xm1d337), a DOE Office of Science User Facility, is supported by the Office of Science of the U.S. Department of Energy operated under Contract No. DE-AC02-05CH11231. This research was supported by the German Research Foundation (DFG) Emmy Noether grant RA 3117/1-1 (to M.T.R.) and seed funding of the CRC1101 “Molecular Coding of Specificity in Plant Processes” (to M.T.R).

## AUTHORS CONTRIBUTION

T.D.G.N. and M.T.R conceived and designed the research. M.T.R. performed and analyzed the RNA-seq data. T.D.G.N., L.S.B., M.W.S, D.Z., L.R., B.J. H.L. and M.T.R. performed the experiments. R.S. andJ.P.V and D.L.C. generated and provided NaN mutant lines. TD.G.N., L.B., M.W.S., D.Z., H.L. and M.T.R. analyzed, interpreted and visualized the data. TD.G.N., H.L. and M.TR. wrote the manuscript. All authors commented on and edited the manuscript.

## DATA AND CODE AVAILABILITY

RNA-seq data have been deposited at GEO and are publicly available (GEO accession number GSE206682). All quantitative data generated and analyzed in this study can be found in the Supplementary Dataset. Microscopy images reported in this paper will be shared by the lead contact upon request. This paper does not report original code. Any additional information required to reanalyze the data reported in this paper is available from the lead contact upon request.

## SUPPLEMENTARY INFORMATION

Supplemental Information includes four figures, three movies, and one supplemental dataset file and can be found online at https://figshare.com/projects/Supplementary_Information_for_Nunes_et_al_2023_-_BdPOX/163402

## MATERIAL AND METHODS

### Plant Material and Growth Conditions

The *B. distachyon* line Bd21-3 was used for all experiments. The *bdpox* mutant lines (*bdpox-1*, NaN1508 and *bdpox-2*, NaN1528) were obtained from the Sodium Azide (NaN_3_) mutagenized population (NaN lines) that was fully resequenced. NaN1508 contained a heterozygous early STOP mutation. Homozygous lines (*bdpox-1*, NaN1508^*bdpox-1*^) and wild-type-like lines (NaN1508^*BdPOX*^) were selected by PCR amplifying the variant containing region using priTN88/priTN89 followed by Sanger sequencing.

Seeds were sterilized for 15 min with 20% bleach and 0.1% Tween, thoroughly rinsed, stratified on MS plates (½ MS (Caisson Labs), 1% Agar (w/v), pH 5.7) for 2 days at 4°C before transfer to a 28°C growth chamber with 16h light:8h dark cycle (110 μmol photons m^-2^ s^-1^) or directly transferred to soil ^67^.

Growth conditions for *B. distachyon* are specified in Nunes et al. (2022) ^7^. In short, plants on soil were grown in a greenhouse with 18h light:6h dark cycle (200-400 μmol m^-2^ s^-1^; day temperature = 28°C, night temperature = 22°C).

### Transcriptional profiling of WT and *bdmute* leaves by RNA-sequencing

25 mature leaf zones (25-30 mm from the base of 2^nd^ leaves) per replicate were collected from three wild-type (Bd21-3) replicates and from three *bdmute* replicates (7 days after germination seedlings grown on ½ MS plates at 20°C with ~100 μmol photons m^-2^ s^-1^ light) were carefully collected, snap-frozen in liquid nitrogen and grounded using mortar and pistil. RNA extraction, library preparation, RNA-sequencing and data analysis was essentially performed as described in Zhang et al. (2022) ^8^. To be specific, total RNA was isolated using Qiagen’s RNeasy Plant Mini kit with on-column DNAse digestion according to the manufacturer’s instructions. The Kapa mRNA HyperPrep (Roche) was used to generate an mRNA enriched sequencing library with an input of 1μg of total RNA. The libraries were sequenced using the Illumina NextSeq500 platform. Read quality was assessed with FastQC and mapped against the Bd21-3v1.0 genome using bowtie2. Mapped reads were counted using summarized overlap and differentially expressed genes were analyzed using DeSeq2. Finally, gene expression was normalized by transcripts per kilobase million (TPM). Raw and processed data are available at Gene Expression Omnibus (GEO) with the accession number GSE206682.

### Reporter Constructs

Reporter and overexpression constructs were generated using the Greengate cloning system ^63^. *BdPOX* promoter and coding sequences were amplified from wild-type *Brachypodium distachyon* (Bd21-3) genomic DNA extracted using a standard CTAB DNA extraction protocol ^68^ and from cDNA synthesized with the RevertAid First Strand cDNA Synthesis Kit (Cat. No.: K1621); ThermoFisher Scientific, Massachusetts, USA) from RNA extracted with the RNeasy Micro Kit (Cat. No.: 74004; Qiagen, Hilden, Germany).

To clone the *BdPOX* genomic sequence, a point mutation was induced in the genomic *BdPOX* sequence to eliminate a BsaI/Eco31I site (GGTCTC) in the second intron. Genomic *BdPOX* was amplified in two separate fragments using priTN99/priTN102 and priTN100/priTN101 (containing the bp substitution **A**GTCTC). The two resulting PCR products were purified using the NucleoSpin Gel and PCR Clean-up kit (Ref. REF 740609.50; Macherey-Nagel, Düren, Germany), digested at 37°C overnight using FastDigest Eco31I (Thermo Fisher Scientific, Waltham, Massachusetts, USA) and ligated overnight at 16°C using T4 ligase (NEB, Ipswich, Massachusetts, USA). The fully reassembled *BdPOX* gene was ligated overnight at 16°C with the previously digested (FastDigest Eco31I) and dephosphorylated pGGC000 entry vector (with Antarctic Phosphatase, NEB) to generate *pGGC_BdPOX. BdPOX (CDS*) sequence was amplified using priTN99/priTN100 from cDNA. The resulting PCR product was purified and digested using FastDigest Eco31I (Thermo Fisher Scientific), and ligated using T4 ligase (NEB) with previously digested (FastDigest Eco31I) and dephosphorylated (Antarctic Phosphatase) pGGC000 entry vector to generate *pGGC_BdPOX(CDS*).

To clone the *BdPOX* promoter, a 3.5kb region upstream of the *BdPOX* transcriptional start site was amplified using priTN95/priTN96. The PCR product was purified and digested using FastDigest Eco31I (Thermo Fisher Scientific) and ligated using T4 ligase (NEB) with previously digested (FastDigest Eco31I) and dephosphorylated (Antarctic Phosphatase) pGGA000 entry vector to generate *pGGA_BdPOXp*. The constitutive *ZmUbi* promoter was amplified from *pEX_ BdMUTEp:CitYFP-BdMUTE ^3^* with priTN3/priTN4 and cloned into pGGA000 (*pGGA_ZmUbip*). The *BdMUTE* promoter was amplified from *pEX_BdMUTEp:CitYFP-BdMUTE* ^3^ with priTN1/priTN28 and priTN2/priTN29 (containing a bp substitution GA**T**ACC to mutate a BsaI site) and cloned into pGGA000 (*pGGA_BdMUTEp*). The PvUbip-driven hygromycin resistance cassette was amplified from *pTRANS_250d* ^65^ using priTN11/priNT30 and priTN12/ priTN29 (containing a 1 bp substitution GA**T**ACC to mutate a BsaI in the *PvUbi2* promoter sequence), and cloned into pGGF000 (*pGGF_PvUBi2p:HygR*). All constructs were test digested and Sanger sequenced to verify that the appropriate sequence was yielded.

The entry modules pGGB003 (B-dummy), pGGD002 (D-dummy) and pGGE001 (rbcS terminator) are described in ^63^. The entry modules *pGGB_3xNLS* and *pGGC_eGFP* were generously provided by Prof. Dr. Karin Schumacher’s group. The pGGD009 (Linker-mCitrine) was generously provided by Prof. Dr. Jan Lohmann’s group.

*BdPOXp:BdPOX-mCitrine, ZmUbip:BdPOX:mVenus, ZmUbi-p:BdPOX(CDS)-mCitrine, BdMUTEp:BdPOX(CDS)-mCitrine* and *BdPOXp:3xNLS-eGFP* were generated using the Green Gate assembly ^63^. In short, the 6 different entry modules (pGGA_ specific promoter; pGGB_dummy or N-tag; pGGC_gene sequence or tag; pGGD_dummy or C-tag; pGGE_terminator and pGGF_resistance) were repeatedly digested and ligated with the destination vector pGGZ004 during 50 cycles (5 min 37°C followed by 5 min 16°C) followed by 5 min at 50°C and 5 min at 80°C for heat inactivation of the enzymes. All final constructs were test digested and the generated GreenGate overhangs were Sanger sequenced.

### Generation and Analysis of Transgenic Lines

Embryonic calli derived from Bd21-3 and *bdpox-1* parental plants were transformed with AGL1 *Agrobacterium tumefaciens* containing the binary expression vectors, selected based on hygromycin resistance, and regenerated as described in Zhang et al. (2022) ^8^. In short, young, transparent embryos were isolated and grown for three weeks on callus induction media (CIM; per L: 4.43g Linsmaier & Skoog basal media (LS; Duchefa #L0230), 30g sucrose, 600μl CuSO4 (1mg/ ml, Sigma/Merck #C3036), 500μl 2,4-D (5mg/ml in 1M KOH, Sigma/Merck #D7299), pH 5.8, plus 2.1g of Phytagel (Sigma/Merck #P8169)). After three weeks of incubation at 28°C in the dark, crisp, yellow callus pieces were subcultured to fresh CIM plates and incubated for two more weeks at 28°C in the dark. After two weeks, calli were broken down to 2-5mm small pieces and subcultured for one more week at 28°C in the dark. For transformation, AGL1 Agrobacteria with the desired construct were dissolved in liquid CIM media (same media as above without the phytagel) with freshly added 2,4-D (2.5μg/ml final conc.), Acetosyringone (200μM final conc., Sigma/Merck #D134406), and Synperonic PE/ F68 (0.1% final conc., Sigma/Merck #81112). The OD600 of the Agrobacteria solution was adjusted to 0.6. Around 100 calli were incubated for at least 10 min in the Agrobacteria solution, dried off on sterile filter paper and incubated for three days at room temperature in the dark. After three days, transformed calli were moved to selection media (CIM + Hygromycin (40μg/ ml final conc., Roche #10843555001) + Timentin (200μg/ ml final conc., Ticarcillin 2NA & Clavulanate Potassium from Duchefa #T0190)) and incubated for one week at 28°C in the dark. After one week, calli were moved to fresh selection plates and incubated for two more weeks at 28°C in the dark. Next, calli were moved to callus regeneration media (CRM; per L: 4.43g of LS, 30g maltose (Sigma/Merck #M5885), 600μl CuSO4 (1mg/ml), pH 5.8, plus 2.1g of Phytagel). After autoclaving, cool down and add Timentin (200μg/ml final conc.), Hygromycin (40μg/ml final conc.), and sterile Kinetin solution (0.2μg/ml final conc., Sigma/Merck #K3253). Calli were incubated at 28°C and a 16h light:8h dark cycle (70-80 μmol PAR m^-2^ s^-1^). After 1-6 weeks in the light, shoots will form. Move shoots that are longer than 1cm and ideally have two or more leaves, to rooting cups (Duchefa #S1686) containing rooting media (per L: 4.3g Murashige & Skoog including vitamins (Duchefa #M0222), 30g sucrose, adjust pH to 5.8, add 2.1g Phytagel). After autoclaving, cool down and add Timentin (200μg/ml final concentration). Once roots have formed, plantlets can be moved to soil (consisting of 4 parts ED CL73 (Einheitserde) and 1 part Vermiculite) and grown in a greenhouse with 18h light:6h dark cycle (250-350 μmol PAR m^-2^ s^-1^). Ideally, the transgenic plantlets moved to soil are initially kept at lower temperatures (day temperature = 22°C, night temperature = 18-20°C) for 2-4 weeks until they have rooted sufficiently before being moved to the warmer greenhouse (day temperature = 28°C, night temperature = 22°C).

We refer to *Brachypodium* regenerants as T0 and to the first segregating population as T1. We analyzed 5-10 independent lines in the T0 generation (depending on how many independent lines were recovered upon regeneration). We confirmed the observed expression pattern and performed the phenotyping studies using one to three independent and fertile T0 transgenics that produced seeds (3-4 T1 individuals per line).

### Gas Exchange Phenotyping

Infra-red gas analyser-based leaf-level gas exchange measurements were performed as described in Nunes et al. (2022) ^7^. All measurements were performed on the youngest fully expanded mature leaf of *B. distachyon* plants 3 weeks after sowing (17-24 days after sowing) using a LI-6800 (LI-COR Biosciences Inc, Lincoln, NE, USA). Ambient light intensity was monitored during the measurements using an external LI-190R PAR Sensor attached to LI-6800. Greenhouse temperature and relative humidity were monitored during the experiments using an Onset HOBO U12-O12 4-channel data logger that was placed next to the plants used for analysis.

#### Light response kinetics

LI-6800 chamber conditions were as follows: flow rate, 500 μmol s^-1^; fan speed, 10000 rpm; leaf temperature, 28° C; relative humidity (RH), 40 %; [CO_2_], 400 μmol mol^-1^; photosynthetic active radiation (PAR), 1000 – 100 – 1000 – 0 μmol PAR m^-2^ s^-1^ (20 min per light step). Gas exchange measurements were automatically logged every minute. The leaf section measured inside the LI-6800 chamber was collected, fixed and cleared to measure leaf area and accurately determine *A* and *g*_sw_ and stomatal anatomical parameters like stomatal length and density. iWUE was calculated as the ratio of *A* to *g*_sw_. Stomatal opening and closure speed were evaluated by rate constants (k, min^-1^) determined from exponential equations fitted for each of the three light transitions (1000-100, 100-1000 and 1000-0 PAR), as described in Nunes et al. (2022) ^7^.

#### Maximum stomatal conductance

Maximum stomatal conductance (physiological *g*_s_max) measurements were performed with the following LI-6800 conditions: flow rate, 500 μmol s^-1^; fan speed, 10000 rpm; leaf temperature, 28°C; relative humidity (RH), 68-70 %; [CO_2_], 100 μmol mol^-1^; PAR, 1500 μmol PAR m^-2^ s^-1^. Gas exchange measurements were automatically logged every minute and physiological *g*_s_max was calculated as the average of the last 5 min at steady-state.

#### Anatomical *g*_s_max calculations

Anatomical *g*_s_max calculations were performed on the 6 individuals for which gas exchange and stomatal anatomy was assessed (Fig. 1C-H). Based on the formula optimized for *B. distachyon* in Nunes et al. (2022) ^7^ this requires the measurement of three anatomical parameters: stomatal length, stomatal density and GC width at the apex (average of 30 stomata per individual) from images obtained using a Leica DM5000B microscope.

#### Steady-state stomatal conductance

LI-6800 chamber conditions were as follows: flow rate, 500 μmol s^-1^; fan speed, 10000 rpm; leaf temperature, 28°C; relative humidity (RH), 40 %; [CO_2_], 400 μmol mol^-1^; photosynthetic active radiation (PAR), 1000 PAR m^-2^ s^-1^ (20 min). Gas exchange measurements were automatically logged every minute. The leaf section measured inside the Li-6800 chamber was collected to measure leaf area to accurately determine *A* and *g*_sw_ and, then fixed and cleared to determine stomatal anatomical parameters like stomatal length and density. All measurements were performed in a semi-randomized manner between 11:30 and 17:30 h to assure measurements for each genotype covered identical periods of time of the day and to avoid the influence of the diurnal variation of *g*_sw_ observed in Nunes et al. (2022) ^7^.

#### Ambient-adapted stomatal conductance

Steady-state ambient-adapted stomatal conductance was assessed using a SC-1 porometer (Meter, Pullman, Washington, USA). SC-1 was calibrated using the calibration plate and the moist circular filter paper provided with the SC-1. Each measurement was performed in automode (30 s). The relative humidity of the SC-1 porometer was returned to < 10 % after each measurement by shaking the sensor head for 30-90 s. Three to four fully expanded leaves per individual were measured twice. Three *B. distachyon* (Bd21-3) and three *bdpox-1* individuals were assessed three weeks after sowing. All measurements were performed in a randomized manner between 8:00 and 9:30 h, to avoid the influence of the diurnal variation of *g*_sw_ observed in Nunes et al. (2022) ^7^.

Important note: Some of the wild-type gas exchange measurements were previously published in Nunes et al. (2022) ^7^, where 120 wild-type measurements over the course of 2 years were correlated to variable growth conditions. The WT gas exchange data that was published in Nunes et al. (2022) ^7^ is indicated accordingly in the Supplementary Dataset.

### Microscopy and Phenotypic Analysis

Most of the morphometric and the cell wall measurements were performed on the actual leaf segments used for gas exchange measurement to thoroughly link cellular form and composition to stomatal gas exchange.

#### Leaf epidermis morphometry

For DIC imaging, the youngest fully expanded mature leaves (3 weeks after sowing) were collected after LI-6800 measurements and placed into 7:1 ethanol:acetic acid and incubated overnight to fix the leaf tissue and remove chlorophyll. To prepare samples for imaging, the tissue was rinsed twice in water, mounted on slides in Hoyer’s solution ^69^ and the abaxial side was examined using a Leica DM5000B microscope (Leica Microsystems, Wetzlar, Germany). Typically, 4-6 (40x objective) and 3-5 (20x objective) abaxial fields of view per leaf of each individual plant were imaged to determine stomatal length, stomatal density, stomatal width at the apices, pavement cell length, prickle hair cell (PHC) base length, base area and/or PHC outgrowth using the straight line tool xyz and/or the polygon selection tool in Fiji ^66^. In the case of complementation experiments represented in Figure 2, the slides were prepared and randomized by an independent researcher before measurements on Fiji, to avoid potential biased phenotyping. The confocal morphometrical analysis of stomata in Bd21-3 and *bdpox-1* mutants, were performed as described in Nunes et al. (2022) ^7^. Leaves were incubated overnight in buffer solution (50mM KCl, 10mM MES-KOH) with 4 mM Fusicoccin (Santa Cruz Biotechnology, Inc., Dallas, TX, USA; Cat. no. 20108-30-9) in the light to force stomatal opening. Leaves were stained in propidium iodide (Sigma-Aldrich, St. Louis, Missouri, USA, Cat. no. P3566; PI, 1:100 of a 1 mg/ml stock) for 5 min and Z-stacks were taken using confocal microscopy. Image analysis was done using Fiji to measure stomatal pore length and stomatal pore area (hand-traced).

#### Stomatal associated-epidermal defects

Stomata and stomata surrounded by defective cell divisions were counted in 5 abaxial fields of view (40 x objective) per leaf of each individual plant. Finally the percentage of stomatal-associated defects was calculated as the total number of stomata surrounded by defective cell divisions using the following formula: sum of 5-7 fields of view)/total number of stomata (sum of 5-7 fields of view) x 100.

#### Reporter lines

For confocal imaging, emerging 2^nd^ (6-7 days post germination (dpg)) or 3^rd^ (11-12 dpg) leaves from plants grown on plates were carefully pulled from the sheath of the older leaf to isolate and reveal the developmental leaf zone. Samples were stained in propidium iodide (PI, 1:100 of a 1 mg/ml stock) for 5 min to stain cell walls and/or mounted directly in water for imaging on a Leica SP8 confocal microscope (Leica Microsystems, Wetzlar, Germany). Image analysis was done using Fiji.

#### Total phenolics autofluorescence

Small leaf fragments previously fixed and cleared in 7:1 ethanol:acetic acid were washed in 70 % ethanol and then transferred to distilled water with 0.02 % (v/v) Tween for rehydration for 3 hours and mounted in distilled water for imaging. Samples were imaged on a Leica SP8 confocal microscope. Excitation and detection settings were as follows: Ex. 405 nm and Em. 490-550 nm. Laser power set to 10 %. For the analysis of PHCs, stacks of 0.33 μm steps were obtained and plot profile analysis was performed on sum slices Z-projections in Fiji. For the analysis of PHCs plot profiles, a straight line was drawn from the base of the PHCs to the tip and plot profiles of gray values were obtained. For the analysis of PHC average autofluorescence, PHCs were handtraced on the sum slices Z-projections and CTCF (corrected total cell fluorescence) calculated as Integrated Density – (Area of selected cell x Mean fluorescence of background readings). Mean fluorescence of 9 background readings per image were obtained to calculate CTCF (using the traced area at regions with only background signal) and to correct plot profiles. For the analysis of mature stomata autofluorescence, GCs were handtraced and autofluorescence was calculated as CTCF (corrected total cell fluorescence) = Integrated Density – (Area of selected cell x Mean fluorescence of background readings) from single images. Mean fluorescence of 3-4 background readings per image were obtained to calculate CTCF (using the traced area at regions with only background signal).

For the analysis of phenolic compounds during stomatal elongation (step 6i-6iii), the emerging 2^nd^ (6-7 dpg) or 3^rd^ (11-12 dpg) leaves were carefully pulled from the sheath of the older leaf to isolate and reveal the developmental leaf zone were mounted in water for imaging on a Leica SP8 confocal microscope. Autofluorescence intensity was measured on handtraced GCs from sum slices Z-projections (75 stacks) using Fiji and calculated as CTCF (corrected total cell fluorescence) = Integrated Density – (Area of selected cell x Mean fluorescence of background readings). Mean fluorescence of 3-4 background readings surrounding each stoma were obtained to calculate CTCF for each GC pair.

#### Basic fuchsin staining

Small leaf fragments previously fixed and cleared in 7:1 ethanol:acetic acid were transferred to distilled water with 0.02 % (v/v) Tween for rehydration for 3 hours. Samples were incubated in 30 μl of 0.01 % Basic Fuchsin (Sigma-Aldrich, St. Louis, Missouri, USA, Cat. no. 857343) for 5 min and washed twice with 30 μl of 50 % glycerol (v/v) for 5 min (2.5 min per wash step), and mounted in 50 % glycerol. Samples were imaged under Ex. 561 nm and Em. 573-603 nm. Stacks of 0.33 μm steps were obtained. For the analysis of PHCs, sum slices Z-projections were performed. A straight line was drawn from the base of the PHCs until the tip and plot profiles of gray values were obtained. Mean gray value of 9 background readings was obtained to correct each measurement. For the analysis of stomata, GCs were handtraced and fluorescence was calculated as CTCF (corrected total cell fluorescence) = Integrated Density – (Area of selected cell x Mean fluorescence of background readings) from single images. Mean fluorescence of 3-6 background readings was obtained for each image to calculate CTCF.

#### Safranin-O staining

Small leaf fragments previously fixed and cleared in 7:1 ethanol acetic acid were transferred to distilled water for 2 hours. Samples were incubated in 50 μl of 0.2 % Safranin-O (Sigma-Aldrich, St. Louis, Missouri, USA, Cat. no. 84120; v/v in 50 % EtOH) for 10 min, washed with 50 % EtOH for 10 min and hydrated in distilled water for 15 min. Samples were mounted in distilled water and imaged under Ex. 488 nm and Em. 530-560 nm (Channel 1 (C1)) and Ex. 561 nm and Em. 570-600 nm (Channel 2 (C2)). Stacks of 0.33 μm steps were obtained. A straight line was drawn from the base of the PHCs until the tip and plot profiles of gray values for C1 and C2 (same defined ROI) were obtained from the sum slices Z-projections. The ratio between the plot profiles from C2 and C1 was obtained. Additionally, images were analyzed using the Fiji macro developed by Baldacci-Cresp et al. 2020 for calculating a ratio from a generated ratiometric image (C2/C1) ^37^.

#### Statistical Analysis

To test for significant differences between two groups we performed unpaired t-tests. One-way ANOVAs and multiple comparison tests were used when comparing more than two groups. Significance was determined when the p value was lower than 0.05. p values are indicated directly in the graphs and details on each analysis described in the figure legends of the respective graphs. All analyses were performed on GraphPad Prism version 9.1.0, GraphPad Software, San Diego, CA, USA, www.graphpad.com.

