## Supplemental Figures S1 - S4 for "Regulation of hair cell and stomatal size by a hair-cell specific peroxidase in the grass *Brachypodium distachyon*"

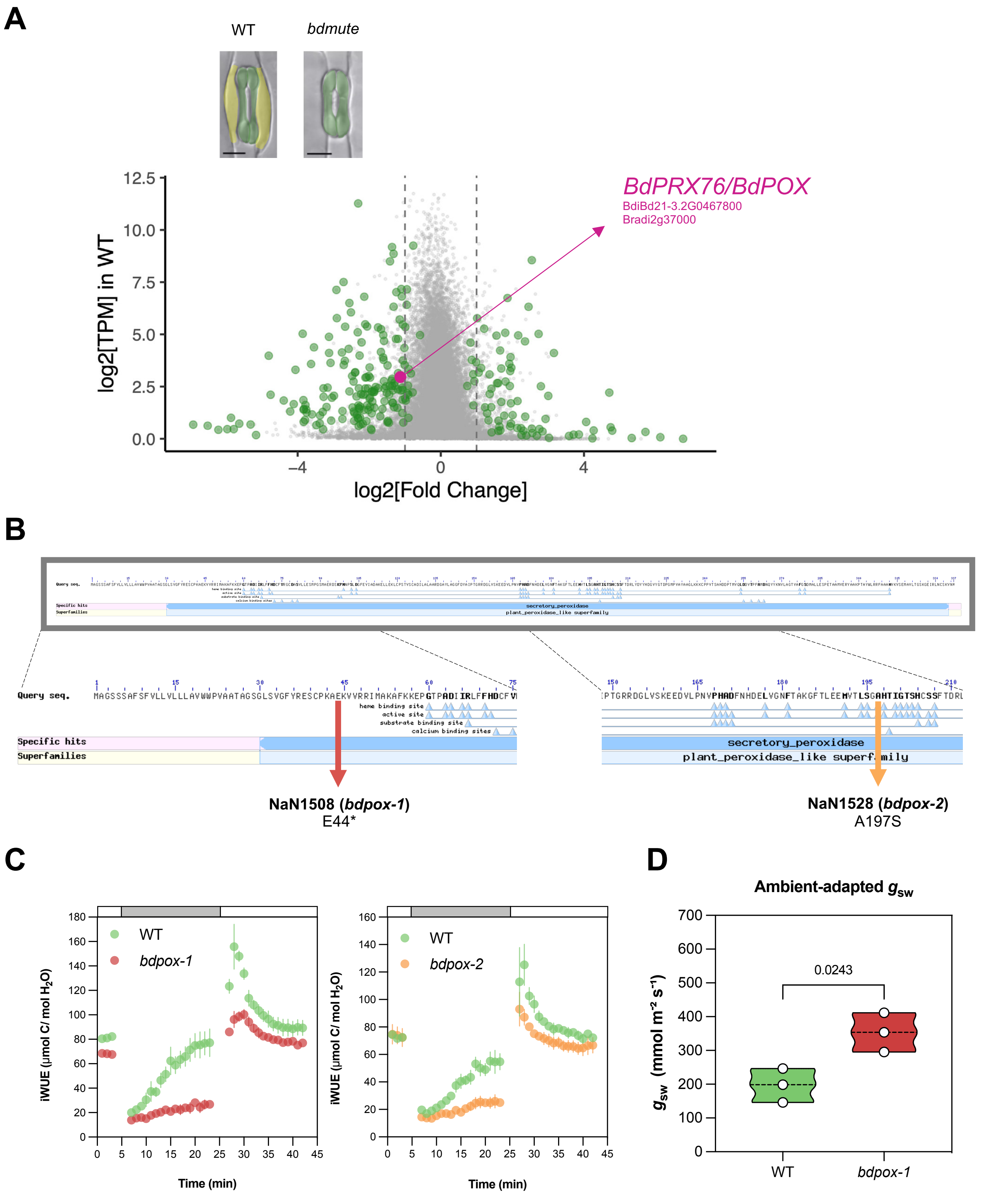


**Figure S1. Comparative RNA-seq and reverse genetic analysis reveals a role for *BdPRX76/BdPOX* in stomatal gas exchange - related to Figure 1. (A)** Volcano plot of RNA-seq data in mature wild type and SC-less *bdmute* leaves. Significantly downregulated genes in *bdmute* are on the left. Green dots represent significantly differentially expressed genes. *BdPRX76/BdPOX* is indicated in magenta. **(B)** Conserved domain analysis of *BdPOX* identified active sites, heme binding site, substrate binding site and calcium binding sites. NaN1508 (*bdpox-1* mutant, in red) contains an early-stop mutation (E44*). NaN1528 (*bdpox-2* mutant, in yellow) contains a non-synonymous mutation in the active/heme binding site (A197S). **(C)** iWUE light-responses (1000-100-1000 PAR) from the initial screening comparing WT and *bdpox-1* (n=3 individuals per genotype), and comparing WT with *bdpox-2* (n=3 individuals per genotype). Error bars represent SEM. **(D)** Ambient-adapted $g_{sw}$ in WT and *bdpox-1* assessed with a SC-1 porometer (n=3 individuals, each point represents the average of two measurements of 3-4 leaves from one individual). P-values obtained from unpaired t-test.

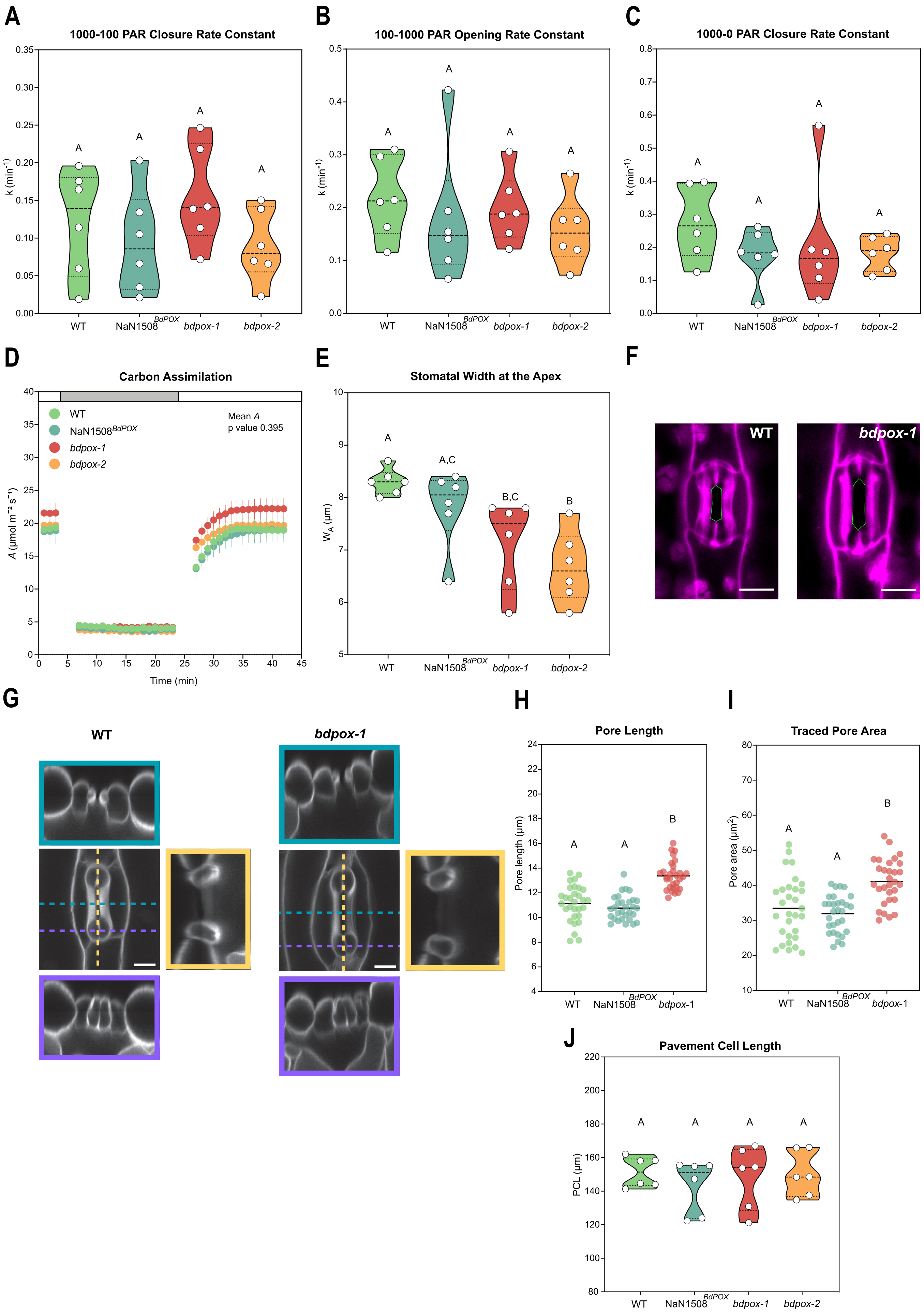
A
1000-100 PAR Closure Rate Constant
B
100-1000 PAR Opening Rate Constant
C
1000-0 PAR Closure Rate Constant
k (min-1)
WT
NaN1508 BdPOX
bdpox-1
bdpox-2
D
Carbon Assimilation
Mean A
p value 0.395
A (μmol m-2 s-1)
Time (min)
E
Stomatal Width at the Apex
WA (μm)
F
WT
bdpox-1
G
WT
bdpox-1
H
Pore Length
Pore length (μm)
I
Traced Pore Area
Pore area (μm2)
J
Pavement Cell Length
PCL (μm)

**Figure S2. Stomatal opening and closure speed, carbon assimilation and detailed stomatal morphometry - related to Figure 1. (A)** Quantification of stomatal closure (1000-100 PAR) rate constant (k) in WT, NaN1508$^{BdPOX}$, *bdpox-1* and *bdpox-2* (n=6 individuals). **(B)** Quantification of stomatal opening (1000-100 PAR) rate constant (k) in WT, NaN1508$^{BdPOX}$, *bdpox-1* and *bdpox-2* (n=6 individuals). **(C)** Quantification of stomatal closure (1000-0 PAR) rate constant (k) in WT, NaN1508$^{BdPOX}$, *bdpox-1* and *bdpox-2* (n=6 individuals). **(D)** Carbon assimilation (*A*) in response to changing light (1000-100-1000 PAR) in WT, NaN1508$^{BdPOX}$, *bdpox-1* and *bdpox-2* (n=6 individuals). **(E)** Stomatal width at the apices ($W_A$) in WT, NaN1508$^{BdPOX}$, *bdpox-1* and *bdpox-2* (n=6 individuals, each point is the average of 30 stomata). **(F)** Open stomatal complexes of WT and *bdpox-1* treated with fusicoccin. Traced stomatal pores highlighted. Scale bars, 10 $\mu$m. **(G)** Orthogonal views of the central section (blue), apex (purple) and the longitudinal axis (yellow) of a wild-type *B. distachyon* stomal complex (left) and a *bdpox-1* mutant stomatal complex (right). Scale bars, 10 $\mu$m. **(H)** Pore length from fusicoccin treated samples in WT, *bdpox-1* and NaN1508$^{BdPOX}$ (n = 30 stomata per genotype). **(I)** Traced stomatal pore area from fusicoccin treated samples in WT, *bdpox-1* and NaN1508$^{BdPOX}$ (n = 30 stomata per genotype). **(J)** Pavement cell length in WT, NaN1508$^{BdPOX}$, *bdpox-1* and *bdpox-2* (n= 6 individuals, 272-301 pavement cells per genotype). Different letters represent significant differences ($p<0.05$). P-values from one-way ANOVA followed by Tukey's multiple comparisons test when more than 3 groups were compared and from unpaired t-tests when comparing 2 groups.

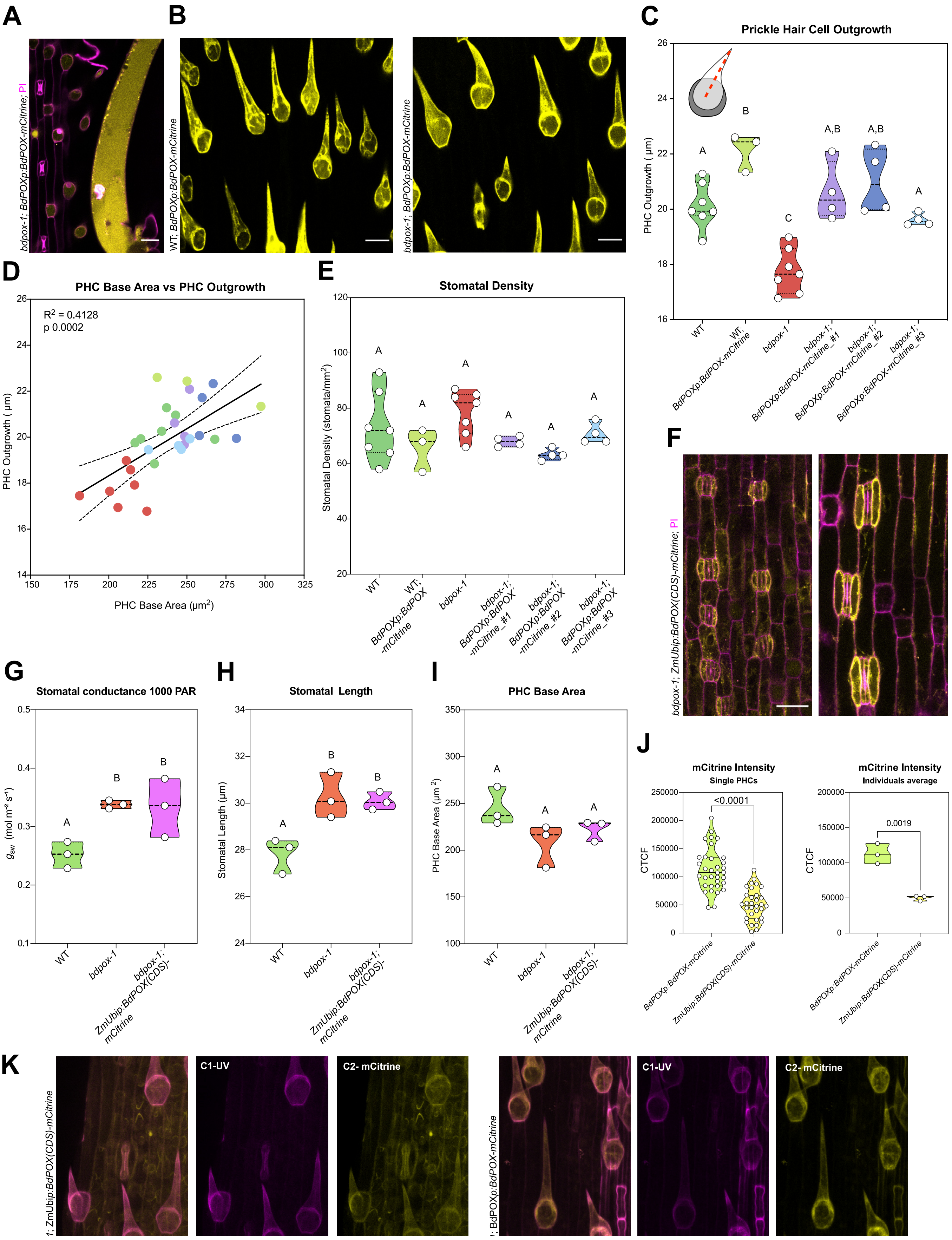
A
B
C
Prickle Hair Cell Outgrowth
PHC Outgrowth (μm)
D
PHC Base Area vs PHC Outgrowth
$R^2$ = 0.4128
p 0.0002
PHC Base Area (μm$^2$)
E
Stomatal Density
Stomatal Density (stomata/mm$^2$)
F
G
Stomatal conductance 1000 PAR
H
Stomatal Length
Stomatal Length (μm)
I
PHC Base Area
PHC Base Area (μm$^2$)
J
mCitrine Intensity
Single PHCs
mCitrine Intensity
Individuals average
CTCF
<0.0001
0.0019
K
C1-UV
C2- mCitrine

**Figure S3. *BdPOX* reporter lines complement the *bdpox-1* phenotype while tissue-wide ectopic expression does not - related to Figure 2.** **(A)** *BdPOXp:BdPOX-mCitrine* expression in prickle hair cells (or microhairs) and in a macrohair. Scale bar, 20 μm. **(B)** *BdPOXp:BdPOX-mCitrine* expression in prickle hair cells in WT and in *bdpox-1*. Scale bar, 20 μm. **(C)** PHC outgrowth in WT, *BdPOXp:BdPOX-mCitrine* (in WT), *bdpox-1* and in three independent complementation lines (*bdpox-1; BdPOXp:BdPOX-mCitrine #1, #2, #3*); n=3-7 individuals per genotype; each dot represents the average of one individual, n= 228-639 PHCs per genotype/line. **(D)** Correlation between PHC base area and PHC outgrowth (n=3-7 individuals per genotype; each dot presents the average traits of one individual). Colors represent the same genotypes as in the other subpanels. **(E)** Stomatal density in WT, WT;*BdPOXp:BdPOX-mCitrine*, *bdpox-1* and in three independent complementation lines (*bdpox-1; BdPOXp:BdPOX-mCitrine #1, #2, #3*); n=3-7 individuals per genotype; 228-639 stomata counted per genotype. **(F)** *ZmUbip:BdPOX(CDS)-mCitrine* expression in the developing leaf (before stomatal elongation) and at mature leaf area (after stomatal elongation and maturation; PI stained cell walls in magenta and mCitrine signal in yellow). Scale bar, 20 μm. **(G)** Steady-state stomatal conductance at 1000 PAR in WT, *bdpox-1* and *bdpox-1; ZmUbip:BdPOX(CDS)-mCitrine* (n=3 individuals per genotype). **(H)** Stomatal length in WT, *bdpox-1* and *bdpox-1; ZmUbip:BdPOX(CDS)-mCitrine* (n= 3 individuals; 133-168 stomata per genotype). **(I)** Prickle hair cell (PHC) base area in WT, *bdpox-1* and *bdpox-1; ZmUbip:BdPOX(CDS)-mCitrine* (n=3 individuals, 129-261 PHCs per genotype). Significant differences represented by different letters ($p<0.05$) obtained from one-way ANOVAs followed by Tukey's multiple comparisons. **(J)** Quantification of mCitrine signal (corrected total cell fluorescence, CTCF) in *bdpox-1; BdPOXp:BdPOX-mCitrine* and *ZmUbip:BdPOX(CDS)-mCitrine* in *bdpox-1;* (n = 3 individuals, 32-34 PHCs per line). **(K)** *ZmUbip:BdPOX(CDS)-mCitrine* and *BdPOXp:BdPOX-mCitrine* expression using the same confocal imaging settings; mCitrine signal in yellow, cell wall UV autofluorescence in magenta. Scale bar, 20 μm.

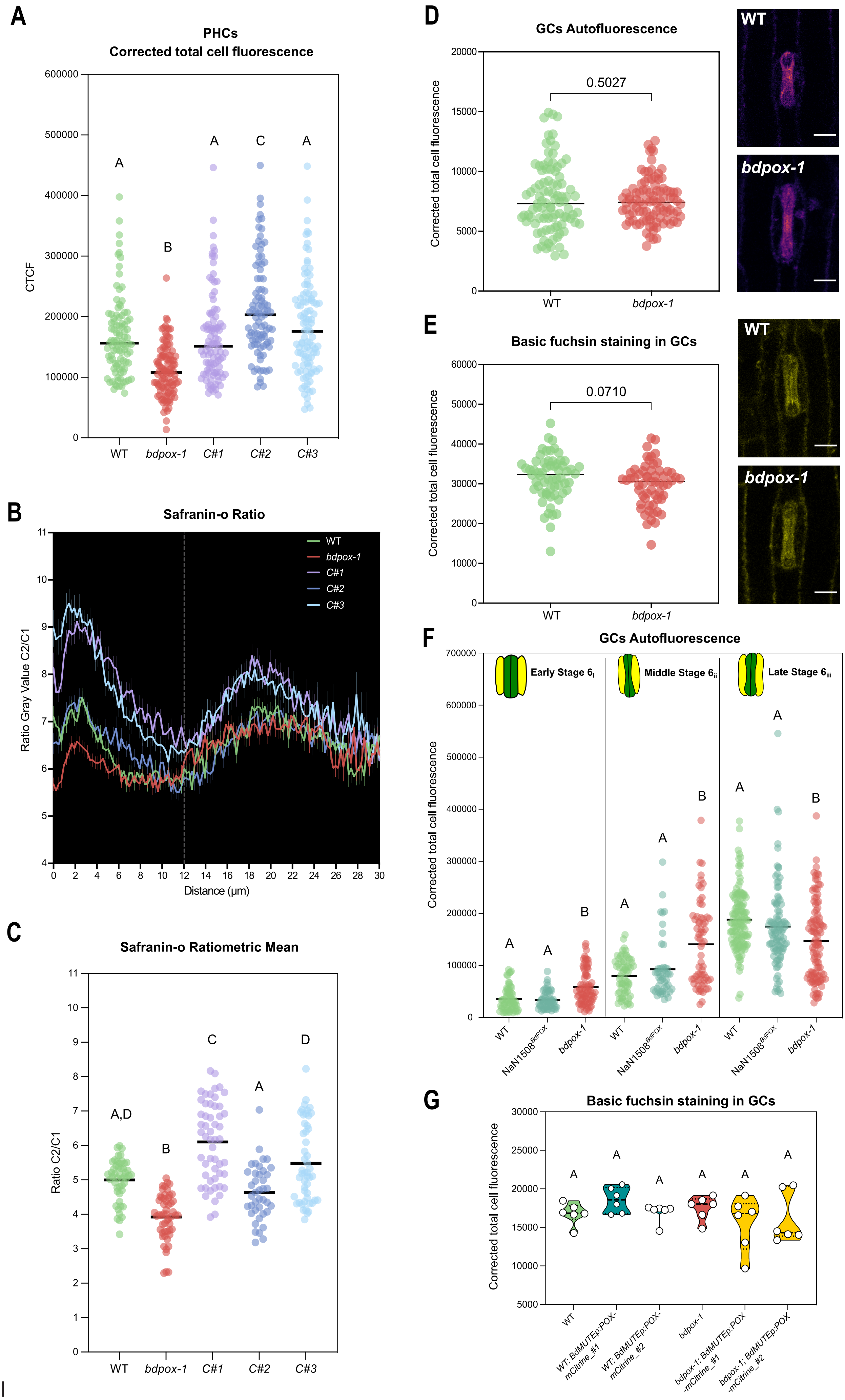
A
PHCs
Corrected total cell fluorescence
CTCF
600000
500000
400000
300000
200000
100000
0
A
B
A
C
A
WT
bdpox-1
C#1
C#2
C#3
B
Safranin-o Ratio
Ratio Gray Value C2/C1
Distance (µm)
WT
bdpox-1
C#1
C#2
C#3
C
Safranin-o Ratiometric Mean
Ratio C2/C1
A,D
B
C
A
D
WT
bdpox-1
C#1
C#2
C#3
D
GCs Autofluorescence
Corrected total cell fluorescence
20000
15000
10000
5000
0
0.5027
WT
bdpox-1
E
Basic fuchsin staining in GCs
Corrected total cell fluorescence
60000
50000
40000
30000
20000
10000
0
0.0710
WT
bdpox-1
F
GCs Autofluorescence
Corrected total cell fluorescence
Early Stage $6_{i}$
Middle Stage $6_{ii}$
Late Stage $6_{iii}$
700000
600000
500000
400000
300000
200000
100000
0
WT
NaN1508$^{BdPOX}$
bdpox-1
G
Basic fuchsin staining in GCs
Corrected total cell fluorescence
30000
25000
20000
15000
10000
5000
WT
WT; BdMUTEp:POX-mCitrine_#1
WT; BdMUTEp:POX-mCitrine_#2
bdpox-1
bdpox-1; BdMUTEp:POX-mCitrine_#1
bdpox-1; BdMUTEp:POX-mCitrine_#2

**Figure S4. Prickle hair cells corrected total cell autofluorescence, stomatal autofluorescence/lignin-staining and additional lignin staining of prickle hair cells - Related to Figure 3 and Figure 4. (A)** Prickle hair cells (PHCs) autofluorescence (corrected total cell fluorescence) in WT, *bdpox-1* and three *bdpox-1;BdPOXp:BdPOX-mCitrine* complementation lines #1, #2 and #3 (n=91-109 PHCs per genotype/line). **(B)** Safranin-o ratio plot profiles in WT, *bdpox-1* and *bdpox-1*; *BdPOXp:BdPOX:mCitrine* lines #1, #2 and #3 (n=45 PHCs per genotype/line). **(C)** Safranin-o ratiometric mean in WT, *bdpox-1* and three *bdpox-1;BdPOXp:BdPOX-mCitrine* complementation lines #1, #2 and #3 (n = 40-53 PHCs per genotype/line). **(D)** GC autofluorescence in mature leaves (3 weeks after sowing) of WT and *bdpox-1* (n=94 - 96 stomata per genotype). Scale bars, 10 $\mu m$. **(E)** GC fuchsin staining in mature leaves (3 weeks after sowing) of WT and *bdpox-1* (n=59 - 62 stomata per genotype). Scale bars, 10 $\mu m$. **(F)** GC autofluorescence during stomatal elongation/maturation ($2^{nd}/3^{rd}$ leaf) in early stage 6i (n=55-78 stomata per genotype/line), middle stage 6ii (n=47-66 stomata per genotype) and late stage 6iii (n=98-129 stomata per genotype) in WT, *bdpox-1* and NaN1508$^{BdPOX}$. **(G)** GC fuchsin staining in mature leaves (3 weeks after sowing) of WT, WT; BdMUTEp:POX-mCitrine #1 and #2, *bdpox-1* and *bdpox-1*; BdMUTEp:POX-mCitrine #1 and #2 (n=6 individuals, 90 stomata per genotype). Scale bars, 10 $\mu m$. Significant differences represented by different letters ($p<0.05$) obtained from one-way ANOVAs followed by Tukey's multiple comparisons.
